# Functional Validation of SAM Riboswitch Element A from *Listeria monocytogenes*

**DOI:** 10.1101/2024.05.28.596223

**Authors:** Ian Hall, Kaitlyn Zablock, Raeleen Sobetski, Chase A. Weidmann, Sarah C. Keane

## Abstract

SreA is one of seven candidate *S*-adenosyl methionine (SAM) class I riboswitches identified in *Listeria monocytogenes*, a saprophyte and opportunistic foodborne pathogen. SAM is essential to all domains of life, serving as a ubiquitous methyl donor and mediator of trans-sulfuration. SreA precedes genes encoding a methionine ATP-binding cassette (ABC) transporter, which imports methionine, a sulfur containing amino acid and substrate for sulfur metabolism. SreA is presumed to regulate transcription of its downstream genes in a SAM-dependent manner. The proposed role of SreA in controlling the transcription of genes encoding an ABC transporter complex may have important implications for how the bacteria senses and responds to the availability of the metabolite SAM in the diverse environments in which *L. monocytogenes* persists. Here we validate SreA as a functional SAM-I riboswitch through ligand binding studies, structure characterization, and transcription termination assays. We determined that SreA has both a similar structure and SAM binding properties to other well characterized SAM-I riboswitches. Interestingly, SreA regulates transcription at a distinctly lower (nM) ligand concentration than other SAM riboswitches but does not substantially terminate transcription, even in the presence of mM SAM.

**GRAPHICAL ABSTRACT:** 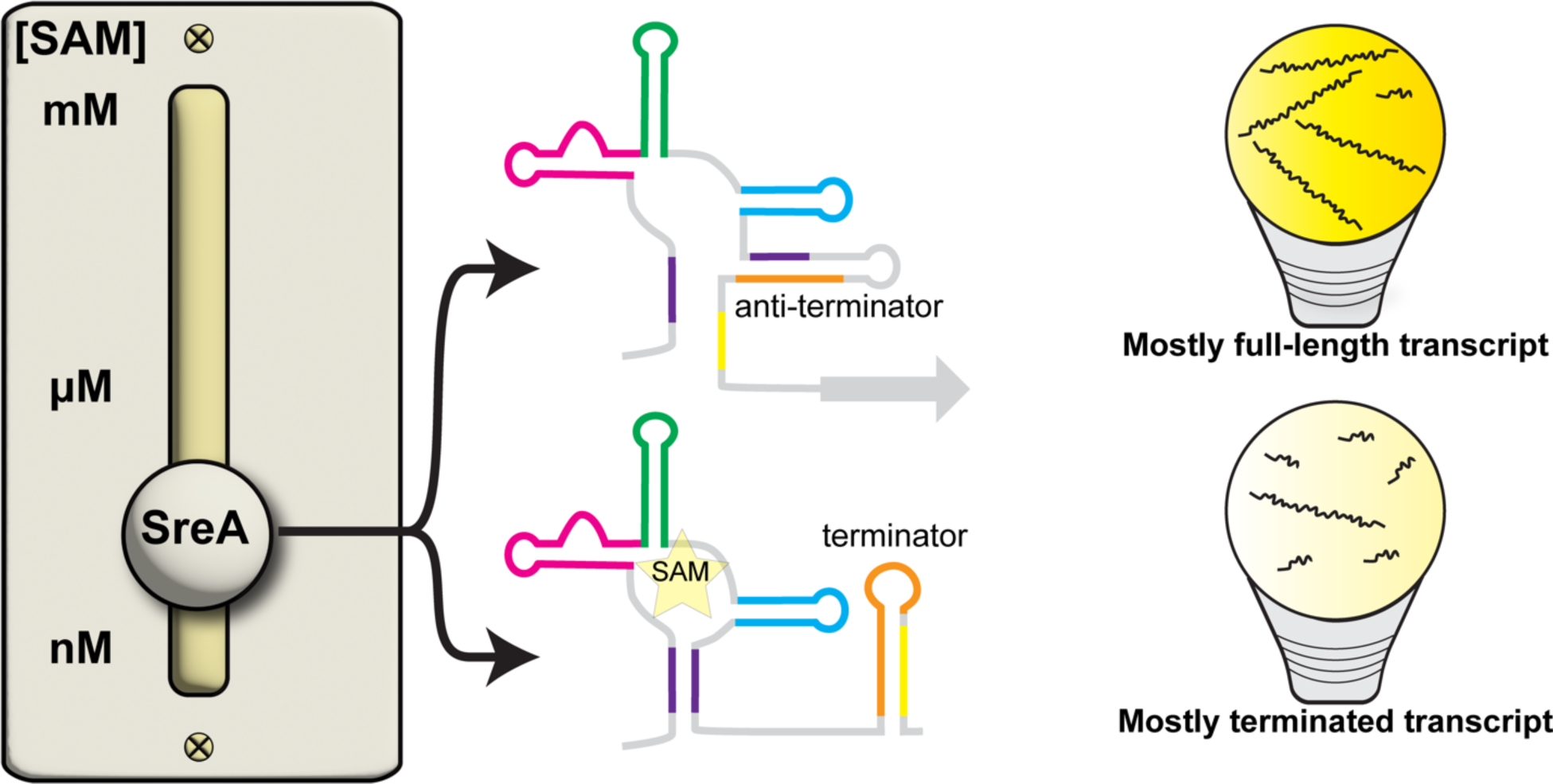

## INTRODUCTION

Riboswitches are structured non-coding (nc) RNAs primarily found in the 5′ untranslated region (UTR) of bacterial messenger (m) RNAs. These RNAs function as switches that can up or down regulate the transcription or translation of the downstream gene in response to binding of a cognate ligand, usually a cellular metabolite or ion.^1–11^ Riboswitches are composed of two functional domains: an aptamer domain, which engages the ligand, and an expression platform, which acts to alter transcription or translation. Ligand binding to an aptamer domain induces conformational changes in the expression platform that either repress translation initiation by sequestration of a ribosome binding site or promote mRNA transcription termination through a rho-independent mechanism.^12^ The genes controlled by a riboswitch typically encode proteins related to the biosynthesis or transport of the riboswitch cognate ligand, creating a regulatory feedback loop.^13^

Translational riboswitches act in a relatively straightforward manner: ligand binding in the aptamer domain inhibits recruitment of the ribosome, whose binding site is typically located within the expression platform. For “OFF” riboswitches, ligand binding stabilizes a riboswitch conformation that sequesters the ribosome binding site through base pairing, thereby repressing translation initiation by blocking small ribosomal subunit recruitment.^12^ Transcriptional riboswitches function in a co-transcriptional manner: only the RNA conformation present while RNA polymerase is transcribing will have a regulatory impact. In many transcriptional riboswitches the folding pathways of a terminator and anti-terminator hairpin compete for shared sequences, and the cognate ligand can alter the folding kinetics of one or the other hairpin to favor one structure in the ensemble (**Fig. 1A**). In the case of an “OFF” riboswitch, ligand binding in the aptamer domain promotes the folding of a specific aptamer domain helix (P1, **Fig. 1A**, right), sequestering a portion of the anti-terminator and enabling the formation of a terminator hairpin with sequence further downstream in the expression platform.^12^ If the RNA polymerase encounters a terminator hairpin, it becomes dislodged from the DNA template and RNA transcript via a rho-independent mechanism thus terminating the transcription of the downstream gene.^12^ In the absence of a ligand, the sequence that would otherwise make up the 3′ half of the P1 stem is instead more readily incorporated into the anti-terminator hairpin in the aptamer domain (**Fig. 1A**, left), and thus RNA polymerase rarely encounters the terminator hairpin during transcription.^12^ The competition between the transcription elongating and terminating states is thought to occur through a ligand-dependent strand displacement mechanism.^14,15^

**Figure 1.**
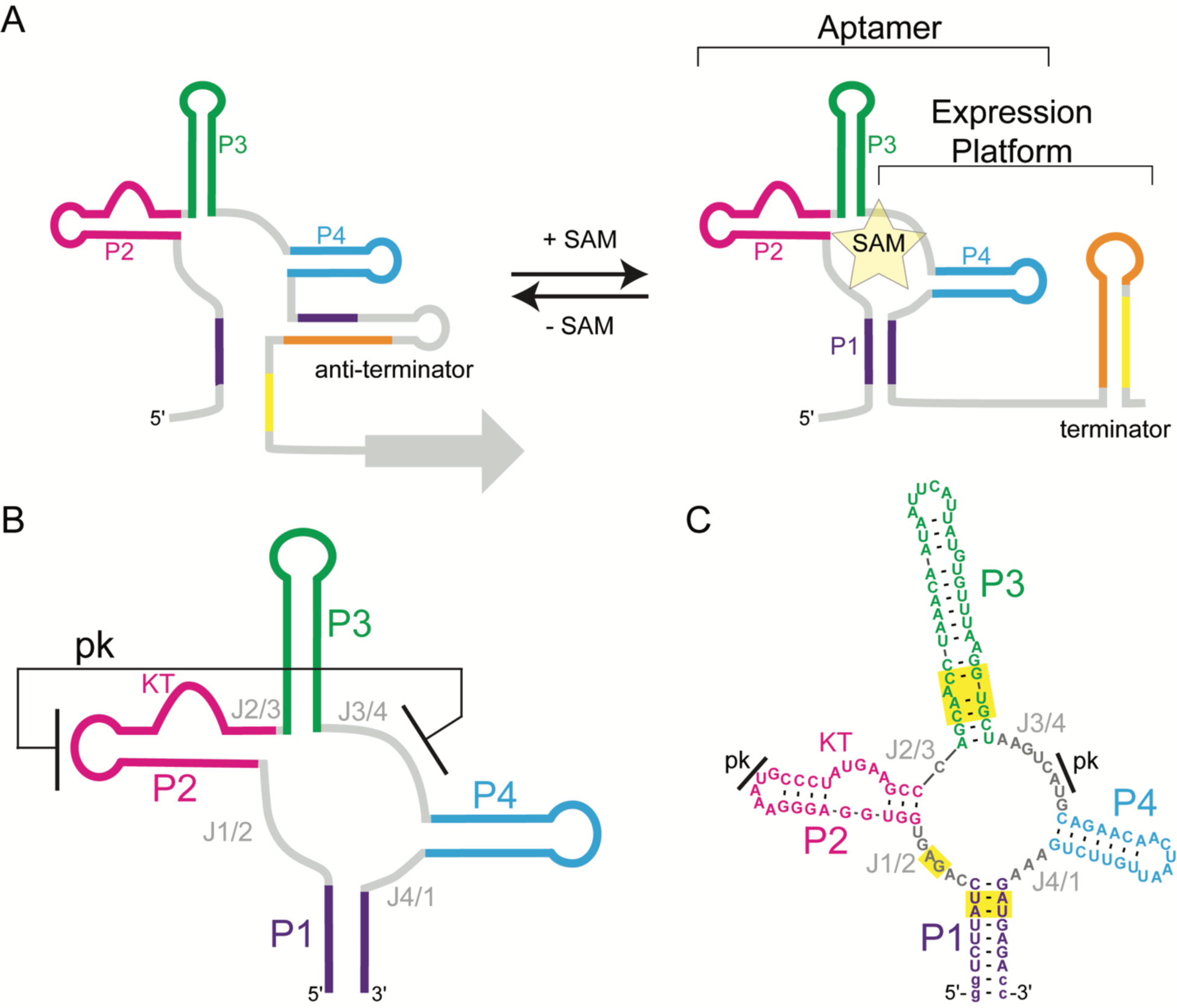
Domain organization of SAM-I riboswitches. **(A)** SAM-I riboswitches sample two major, mutually exclusive conformations during transcription. In the absence of SAM, the anti-terminator structure (left) is favored, and the RNA polymerase transcribes the downstream gene. Upon SAM binding, there is a conformational rearrangement in both the aptamer domain and expression platform and the riboswitch primarily adopts a terminator structure that terminates transcription before the downstream genes are transcribed (right). **(B)** Cartoon of the conserved SAM-I aptamer domain structure. Four helices (P1-P4) and junction regions (J1/2, J2/3, J3/4, and J4/1) are present and there is a pseudoknot (pk) formed between residues in the loop of P2 and J3/4. **(C)** Predicted secondary structure of SreA aptamer domain. Nucleotides predicted to be in a pk are indicated with black lines. Yellow boxes highlight nucleotides expected to be involved in SAM binding. Lowercase nucleotides are non-native.

*S*-adenosylmethionine class-I (SAM-I) riboswitches bind their cognate ligand SAM, a ubiquitous methyl donor in the cell, and regulate the transcription of genes related to sulfur metabolism.^8,16–25^ SAM-I riboswitches bind the cognate ligand with high selectivity over near-cognate ligands like S-adenosylhomocysteine (SAH), a product of SAM methyl transfer reactions.^25,26^ The structures of SAM-I riboswitch aptamer domains from *B. subtilis* and *Thermoanaerobacter tengcongensis* reveal a conserved fold with four helices (P1 – P4) arranged in a four-way junction (J1/2, J2/3, J3/4, J4/1) (**Fig. 1B**).^17,20,21,25,27^ A highly conserved kink-turn motif orients the apical loop of P2 for pseudoknot formation with J3/4 and helps to form a SAM binding pocket in the core of the aptamer domain between P1, P3, and the P1/P2 junction.^17,20,27^

Seven candidate SAM-I riboswitches were identified in the *Listeria monocytogenes* EGD-e genome by tiling arrays and were subsequently named SAM riboswitch elements (Sre) A-G.^28,29^ SreA-G share high sequence similarity with known SAM-I riboswitches and precede genes related to sulfur metabolism. Interestingly, one candidate riboswitch, SreA, was proposed to have a *trans*-acting regulatory function in which it controls the translation of a gene important for virulence in *L. monocytogenes*.^29^ Still, the canonical *cis-*acting SAM-I riboswitch function of SreA has yet to be functionally validated. The predicted secondary structure of SreA has highly conserved SAM binding nucleotides appropriately positioned in P1, J1/2, and P3, suggesting that the RNA may be competent to bind SAM (**Fig. 1C**). In the present work, we experimentally validated SreA as a functional SAM-I riboswitch. We show that SreA binds to SAM with high affinity and that the interaction is specific for SAM and not the near-cognate ligand SAH. Additionally, we characterized the SreA riboswitch structurally using a combination of small-angle X-ray scattering (SAXS) and selective 2′-hydroxyl acylation analyzed by primer extension (SHAPE) and mutational profiling (MaP). Furthermore, we demonstrate that SreA terminates transcription in a SAM-dependent manner. These data collectively validate SreA as a functional SAM-I riboswitch and reveal that SreA is responsive to distinctly low concentrations of SAM, yet is unable to fully terminate transcription of the downstream genes even under mM concentrations of SAM.

## METHODS

### Construct design

The sequence of SreA (**Table S1**) was identified in the *L. monocytogenes* EGD-e genome.^28,29^ A gene block (**Table S2**) containing a T7 promoter, a 5′ hammerhead (HH) ribozyme, the full-length SreA sequence, and flanking restriction sites was ordered from Integrated DNA Technologies (IDT). The gene block was restriction digested and cloned into the pUC57 plasmid. The conserved aptamer domain of SreA was identified by secondary structure prediction using the Dynalign algorithm from RNAstructure.^30,31^ The *B. subtilis yitJ* SAM-I riboswitch sequence (**Table S1**) was used to constrain full-length SreA folding predictions.^19^

### DNA amplification

DNA template for the SreA aptamer domain was generated by PCR amplification of the HH SreA pUC57 plasmid with EconoTaq PLUS 2x Master Mix (Lucigen) using primers that are complementary to the 5′ and 3′ ends of P1 (**Table S3**). The forward primer included a T7 RNA polymerase promoter sequence followed by two guanosine residues to facilitate transcription initiation with T7 RNA polymerase. The reverse primer included two 2′-O-methyl guanosine residues on the 5′ end to help limit non-templated transcription.^32^ These primers result is the addition of two non-native G-C base pairs at the base of the P1 stem.

### RNA transcription and purification

SreA aptamer domain RNA was prepared by *in vitro* transcription as previously described.^33^ Briefly, transcription reactions were carried out in 1x transcription buffer (40 mM Tris-base pH=8.5, 5 mM DTT, 1 mM spermidine, 0.01% Triton-X) using T7 RNA polymerase (prepared in house). The transcription reaction contained 3-6 mM rNTPs, 10-20 mM MgCl_2_, 30-40 ng/μL DNA template, 0.2 U/mL yeast inorganic pyrophosphatase (NEB),^34^ ∼15 µM T7 RNA polymerase, and 10-20% (v/v) dimethyl sulfoxide (DMSO).^35^ Reactions were incubated at 37 °C for 3-4 hours and then quenched using a solution of 7 M urea and 250 mM ethylenediaminetetraacetic acid (EDTA) (pH=8.5). SreA aptamer domain RNA was purified from the reaction by preparative scale denaturing polyacrylamide gel electrophoresis (PAGE). The target RNA was UV shadowed, excised, and electroeluted from the gel in 1x TBE (44.5 mM Tris-base, 44.5 mM boric acid, 5.0 mM EDTA, pH=8.3). The RNA was spin-concentrated, salt-washed with 2 M ultrapure NaCl, and exchanged into water using Amicon Centrifugal Filter Units (MilliporeSigma).

### Isothermal Titration Calorimetry (ITC)

SreA aptamer domain RNA in water was denatured by heating at 95 °C for 3 min and immediately cooled on ice for 3 min. 10x ITC buffer was added the SreA aptamer domain RNA to a final 1x buffer composition of 20 mM HEPES pH=6.5, 5 mM MgCl_2_, 135 mM NaCl. The final RNA concentration was ∼ 30 μM. The sample was allowed to fold in 1x ITC buffer with incubation for 20 min at 37 °C. SAM-Iodide (Sigma) or SAH (Cayman Chemical) was resuspended in water and 10x ITC buffer was added such that the final ligand concentration was ∼300 μM in 1x ITC buffer. The RNA and ligand samples were filtered through a 0.22 μm centrifuge tube filter (Costar) by centrifuging at 16,873 x g for 1 min. ITC experiments were performed in triplicate on a MicroCal PEAQ-ITC calorimeter (Malvern Panalytical) at 25 °C using high feedback mode, a reference power of 10 μcal/s, a stir speed of 750 rpm, and an injection rate of 0.5 μL/s. After an initial delay of 120 s, a 0.2 μL injection was delivered to the sample cell. The initial injection was followed by 19 2 μL injections spaced 160 s apart. Integrated heat data from appropriate control experiments were subtracted using the mean heat method. Integrated heat data were analyzed using a one-site binding model with MicroCal PEAQ-ITC Analysis Software.

### Small angle X-ray scattering (SAXS)

SreA aptamer domain RNA was denatured as described above. Denatured SreA aptamer domain RNA was added into 10x SAXS buffer to a final RNA concentration of 2.5 mg/mL and a final 1x buffer composition of 50 mM potassium phosphate (pH=6.5), 5 mM MgCl_2_, and 135 mM NaCl. The sample was refolded in 1x SAXS buffer with incubation for 20 min at 37 °C. An additional sample was prepared in the same manner except 1 mM SAM was added before refolding. SAXS was performed at BioCAT (beamline 18 ID at the Advanced Photon Source, Chicago) with in-line size exclusion chromatography (SEC) to separate homogenous RNA from aggregates and other contaminants. SreA aptamer domain RNA (200 μL, 2.5 mg/mL) ± 1 mM SAM was loaded onto a Superdex 75 Increase 10/300 GL column (Cytiva) run at 0.6 mL/min and 22 °C by an AKTA PURE FPLC (GE). The eluate was passed through a UV detector and then into the SAXS flow cell, consisting of a 1.0 mm ID quartz capillary with ∼20 μm walls. A coflowing buffer sheath separated samples from the capillary walls, reducing radiation damage.^36^ Scattering intensity was recorded using an Eiger2 XE 9M (Dectris) detector. The detector was placed 3.654 m from the sample giving access to a q-range of 0.0027 Å^-1^ to 0.42 Å^-1^. 0.5 s exposures were acquired every 1 s during elution and data was reduced using BioXTAS RAW 2.1.3.^37^ A buffer blank was created by averaging frames preceding the elution peak and then subtracted from exposures selected from the elution peak to make the I(q) vs q curves used for subsequent analyses. For the (+) SAM sample, small amounts of free RNA (not bound to SAM) were evident in the scattering data and the scattering contributions from the SAM bound RNA were isolated by evolving factor analysis.^38^ The GNOM package was used to determine the pair-distance distribution function [p(r)].^39^ The maximum linear dimension of the molecule, D_max_, was calibrated for goodness-of-fit by enforcing a smooth zeroing of p(D_max_). Full details of SAXS data collection and analysis are presented in the Supporting Information and in **Table S4**.

### Switch-MaP of SreA aptamer domain

Template switching and mutational profiling (Switch-MaP) was used to collect reactivity data for each nucleotide in the SreA aptamer domain.^40^ First, SreA aptamer domain RNA ± 1 mM SAM was reacted with 10 mM 1-methyl-7-nitroisatoic anhydride (1M7) or with DMSO solvent. For both (+) SAM and (-) SAM reactions, 80 pmol of SreA aptamer domain RNA was diluted into 1,120 μL of water and denatured as described above. 320 μL of 5x folding buffer (500 mM HEPES pH=8.0, 500 mM NaCl, 25 mM MgCl_2_) was added to the RNA, and the sample was incubated for 20 min at 37 °C. The RNA sample was split into two 720 μL aliquots. While incubating on a 37 °C heat block, one aliquot was added to 80 μL of anhydrous DMSO (VWR) and the second aliquot was added to 80 μL of 100 mM 1M7 (Sigma) prepared in anhydrous DMSO. The samples were incubated for 2 min at 37 °C and 250 rpm on the heat block in great excess of 1M7’s aqueous half-life of 17 s.^41^ The 1M7 and DMSO-treated samples were exchanged into water using Amicon Centrifugal Filter Units (MilliporeSigma).

After acylation or treatment with DMSO, a 5′ pre-adenylated DNA adapter (**Table S3**) was ligated to the 3′ end of the SreA aptamer domain RNA.The adapter sequence serves as the binding site for reverse transcription initiation and as a platform for downstream library creation.^40^ Briefly, 20 pmol of washed SreA aptamer domain RNA from the 1M7 treated and DMSO treated samples and 60 pmol of the DNA adapter were ligated at 25 °C for 2 h in a reaction mixture containing 10% DMSO, 25% PEG 8000, 10 U T4 RNA ligase 1 (NEB), 50 mM Tris-HCl pH=7.5, 10 mM MgCl_2_, and 1 mM DTT. RNA was purified from the reaction mixture using the RNA Clean and Concentrator-5 kit (Zymo) and eluted in 20 μL of water.

To prepare the RNA for a MaP-RT reaction with a template switching oligonucleotide (TSO, **Table S3**) the Vaccinia Capping System (NEB) was used without the addition of SAM to add a guanylate cap (G_cap_) structure the 5′ end of the adapter ligated SreA aptamer domain RNA. 20 μL of purified RNA from the ligation step was denatured by heating at 65 °C for 5 min and cooling on ice for 5 min. Denatured RNA was incubated at 37 °C for 30 min in a reaction containing 0.5 mM GTP and 0.5 U/μL vaccinia capping enzyme and 1x capping buffer (50 mM Tris-HCl pH=8.0, 5 mM KCl, 1 mM MgCl_2_, 1 mM DTT).

The adapter-ligated, G_cap_ SreA aptamer domain RNA was used as a template in a Switch-MaP-RT reaction. The inclusion of a TSO in the MaP-RT reaction ultimately appends an additional sequencing platform to the 3′ end of the cDNA product.^40^ For the Switch MaP-RT reactions, 2 pmol of adapter-complementary RT primer (**Table S3**) was mixed with 5 pmol of adapter-ligated and 5′-capped SreA aptamer domain RNA. To RNA primer mix, 20 nmol of dNTPs (5 nmol each base) was added (9 µl of total volume of RNA, primers, and dNTPs), heated to 70 °C for 5 min and then immediately placed at 4 °C for 2 min. To this template solution, 9 µl of freshly made 2.22x MaP buffer (111 mM Tris-HCl (pH=8.0), 167 mM KCl, 13.3 mM MnCl_2_, 22 mM DTT and 2.22 M betaine) and 1 µl of 100 μM TSO was added, and the mixture was incubated at 25 °C for 2 min. After adding 200 U of SuperScript II reverse transcriptase (Thermo), reaction mixtures were incubated for 10 min at 25 °C and for 90 min at 42 °C, cycled ten times between 42 °C and 50 °C with each temperature incubation 2 min long and then heated to 70 °C for 10 min to inactivate enzyme.

### Library preparation and sequencing

Targeted amplicon sequencing libraries were generated by PCR. Briefly, 1 µl of complementary DNA (cDNA) from the MaP reverse transcription reaction was used as template for PCR with Q5 hot-start polymerase (NEB) according to manufacturer’s recommendations. PCR products were purified with SPRI beads at a 1x ratio and eluted into 15 µl of nuclease-free water. Size distributions and purities of amplicon libraries were verified (4150 TapeStation, Agilent). About 120 amol of each library was sequenced on a MiSeq instrument (Illumina) with 2 x 150 paired-end sequencing.

### Analysis with SHAPEmapper-2

FASTQ files from sequencing runs along with a FASTA file including the riboswitch sequence were directly input into ShapeMapper 2 software for read alignment to the SreA sequence and mutation counting.^42^ ShapeMapper2 defaults were used apart from the – min-depth flag, which was reduced to 4000 (although all experiments exceeded this threshold at every nucleotide position). The normalized SHAPE reactivities (‘.shape’ output) are required for downstream structural modeling and plotting. RNAstructure Fold with the -mfe flag and the -sh flag (with the .shape output of ShapeMapper2) was used to generate minimum free energy structures from SHAPE data.^30,42^

### Standard score comparison of SAM treated and untreated RNA SHAPE reactivities

SHAPE reactivities of SAM (+) and SAM (-) RNA samples were normalized to each other using a median difference minimization strategy (generalized reduced gradient) to enable sensitivity to single-nucleotide level differences as previously described.^43^ In this case, the log-transformed relative reactivities – derived as log_10_(modified rate/unmodified rate) from the “profile.txt” ShapeMapper2 output – of the SAM (-) samples were scaled down to minimize the median absolute difference between all nucleotides in SAM (+) and SAM (-) samples. Standard scores (Z-scores) were calculated from the resulting set of scaled differences between SAM (+) and SAM (-) relative reactivities (this set of differences is normally distributed).

### Single round *in vitro* transcription termination assays

The template for *in vitro* transcription termination assays was ordered as a gene block (IDT) that contained the T7A1 promoter sequence, a U-deficient sequence that allows the formation of a halted complex, the native riboswitch sequence, and 30 nucleotides of the native downstream gene (**Table S2**).^44^ Single-round transcription reactions were carried out with 4.5 nM template DNA, 9 mU/μL *Escherichia coli* RNA polymerase holoenzyme (NEB), 10 μM rNTPs (Sigma), 0-8 mM SAM (Sigma) and 0.5 mg/mL heparin (Thermo). DNA templates were incubated with *E. coli* RNA polymerase holoenzyme in 1x transcription buffer (20 mM Tris-HCl pH 8.0, 20 mM MgCl_2_, 20 mM NaCl, 100 μM EDTA) at 37 °C for 5 min. AU dinucleotide (100 μM) (Horizon Discovery), ATP (25 μM), CTP (25 μM), and α-^32^P GTP 50 nM (3000 Ci/mmol) (Perkin Elmer/Revvity) were added to the reaction mixture to form a halted complex. The reaction mixture was passed through a G50 spin column (Cytiva) to remove unincorporated AU dinucleotide and NTPs. The purified halted complex was then aliquoted into new tubes containing unlabeled NTPs necessary to transcribe the remainder of the RNA, SAM, and heparin. The addition of heparin concurrently with the NTPs prevents re-initiation of transcription. The reactions were incubated for an additional 15 min at 37 °C. After incubation, the reactions were quenched by adding loading buffer (95% formamide, 1 mM EDTA, 0.1% SDS, 0.2% bromophenol blue, and 0.2% xylene cyanol). Transcription products were resolved by denaturing PAGE on 0.3 mm thick 10% polyacrylamide gels. Phosphor imaging screens (Molecular Dynamics) were exposed to the gels at -70 °C for 16-24 hours. Radiosensitive screens were imaged with an Amersham Typhoon (Cytiva) using phosphor imaging and images were quantified with ImageJ.^45^ The intensity of bands for full length and terminated transcription products was determined by fitting gaussian curves to the reduced intensity vs position data obtained in ImageJ. The half maximum termination response (T_50_) was determined by calculating the proportion of full-length and terminated transcription product at each SAM concentration. The proportion of each product was plotted against the ligand concentration and fit with a sigmoidal four parameter logistic curve where b is the Hill slope (**Equation 1**).

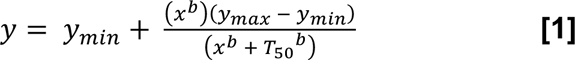

Final reported T_50_ values derive from analysis of the terminated products.

### RNA structure drawing

All RNA structures were rendered using RNAcanvas.^46^

## RESULTS

SreA binds SAM with high affinity and selectivity.

To determine if SreA acts as a canonical SAM-I riboswitch, we first assessed SreA aptamer domain binding to SAM by ITC. The SreA aptamer domain binds SAM spontaneously through an enthalpically driven process (**Fig. 2**, **Table 1**). The energetics of SAM binding to the SreA aptamer domain are consistent with broadly observed enthalpy-driven RNA folding and RNA-ligand interactions.^47–49^ The SreA aptamer domain binds one SAM molecule with a measured K_D_ of 490 ± 34 nM (**Fig. 2**, **Table 1**), suggesting that the isolated aptamer domain is properly folded. The observed SAM binding K_D_ is consistent with that of other SAM-I riboswitches, where K_D_ values for SAM binding range from low nanomolar to low micromolar.^8,16,25,27^

**Figure 2.**
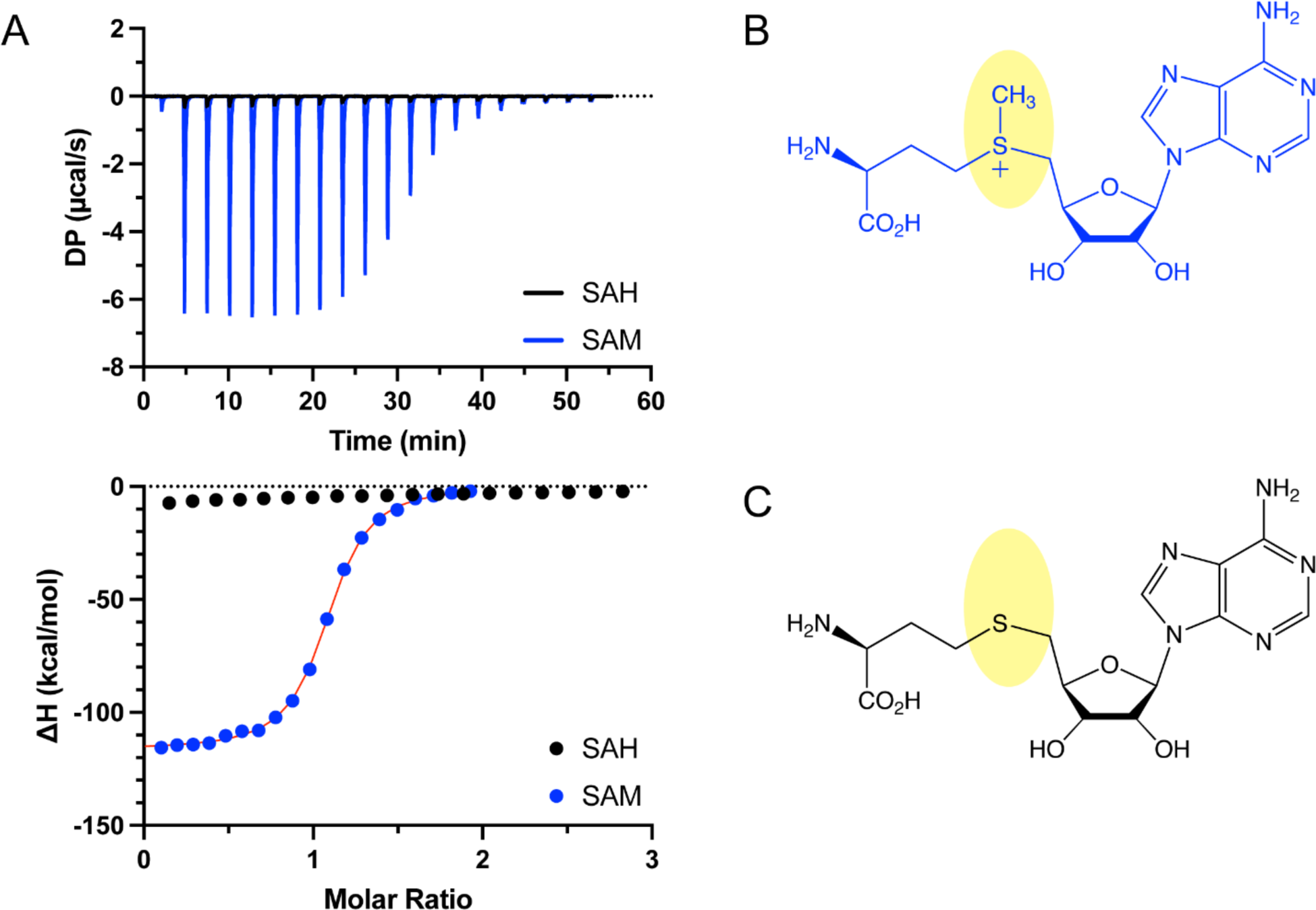
SreA aptamer domain binds SAM but not SAH. **(A)** ITC titration experiments of the SreA aptamer domain binding to SAM (blue) and SAH (black). **(B)** Chemical structure of SAM. **(C)** Chemical structure of SAH. Yellow ovals highlight the minor chemical differences between SAM and SAH.

**Table 1.**
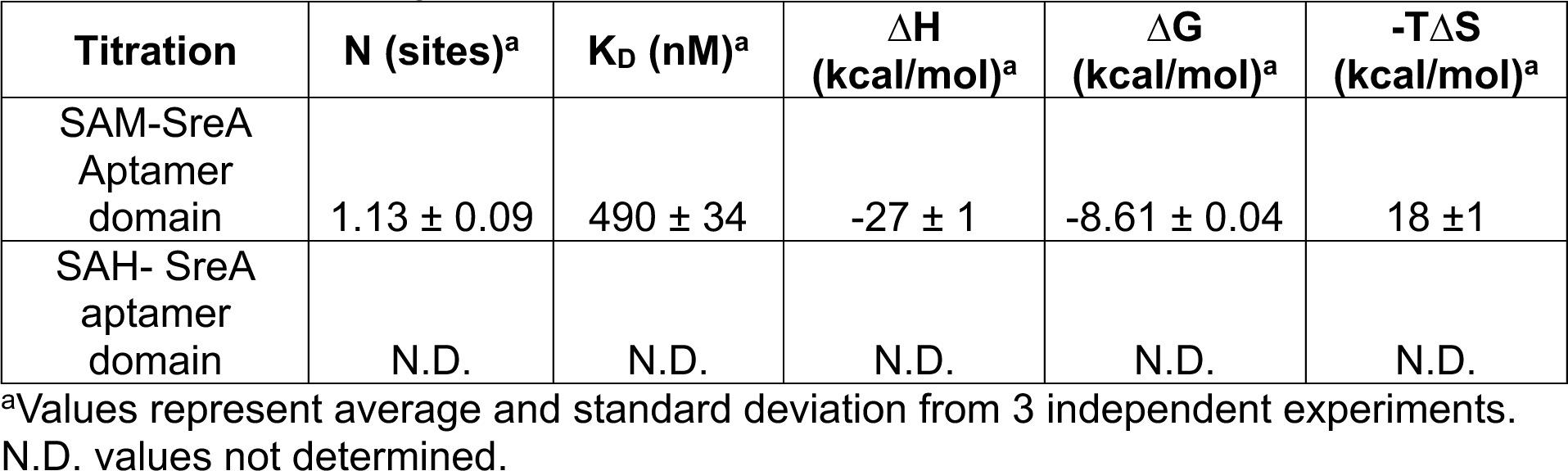
ITC Thermodynamic Data.

Additionally, we show that the SreA aptamer domain binds SAM with high selectivity. Titration of SreA aptamer domain with a near cognate ligand (SAH) resulted is negligible heats of binding, which could not be reliably fit with a one-site binding model (**Fig. 2**). Accordingly, binding to SAH is sufficiently weak relative to that of SAM and not suitable to study by ITC. Consistent with our findings, SAH binding was reportedly ∼ 100-fold weaker for a SAM-I riboswitch from *B. subtilis*.^8^

SreA compacts upon SAM binding.

SAXS data was collected on SreA aptamer domain in the absence and presence of 1 mM SAM (**Fig. 3A,B**). Kratky analysis of the (-) SAM scattering data suggests the SreA aptamer domain RNA is partially folded with flexible regions (**Fig. 3C**). From 0 < q < 0.1, the q^2^*I data resembles a broad Gaussian-like curve suggesting that the SreA aptamer domain is compact. However, from 0.1 < q < 0.2, the q^2^*I data increases to a local maximum and has a q^2^*I plateau at q > 0.2 suggesting that the molecule also has flexible regions. The addition of ligand increases the maximum of the Gaussian-like curve from 0 > q > 0.1 suggesting that ligand binding further compacts the SreA aptamer domain. Additionally, for the (+) SAM sample, q^2^*I trends towards 0 from 0.1 < q < 0.4 and is consistent with a more well-folded structure (**Fig. 3C**). The pair distance distribution function, p(r), analysis of the (-) SAM sample shows that the most probable linear distance between all possible pairs of atoms in the RNA is 40 Å. The addition of ligand shifts the most probable distance to 20 Å, which corresponds to the width of an RNA A-form helix (**Fig. 3D**). This observation supports SAM-dependent stabilization of helical regions within the SreA aptamer domain, as observed in other systems.^21^ The extent of macromolecular compaction upon ligand binding was quantitatively assessed through changes in the radius of gyration (R_g_) and the maximum dimension (D_max_). Upon SAM binding the SreA aptamer domain compacts from an R_g_ of 31.6 to 27.3 and from a D_max_ of 117.0 to 102.0 (**Table 2**). Together the Kratky and p(r) analysis suggests that the SreA aptamer domain binds SAM in a conformationally flexible state and compacts around the ligand.

**Figure 3.**
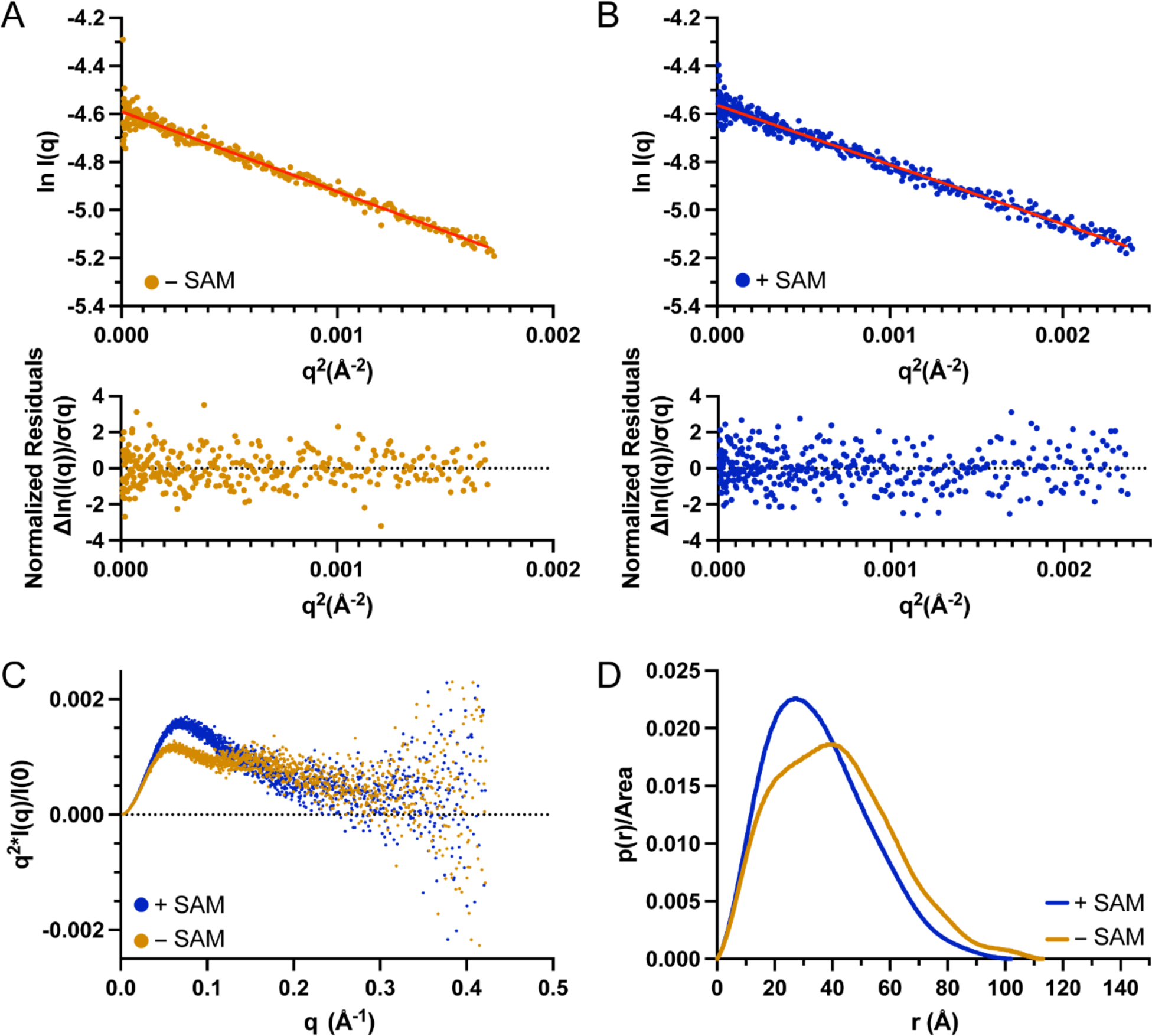
Ligand binding compacts the SreA aptamer domain. **(A)** Guinier fit and normalized residuals for SreA aptamer domain (-) SAM. **(B)** Guinier fit and normalized residuals for SreA (+) 1 mM SAM. **(C)** Dimensionless Kratky plots of SreA aptamer domain (-) SAM (gold) and SreA aptamer domain (+) 1 mM SAM (blue). **(D)** Area normalized p(r) plots of SreA aptamer domain (-) SAM (gold) and SreA aptamer domain (+) 1 mM SAM (blue).

**Table 2.**
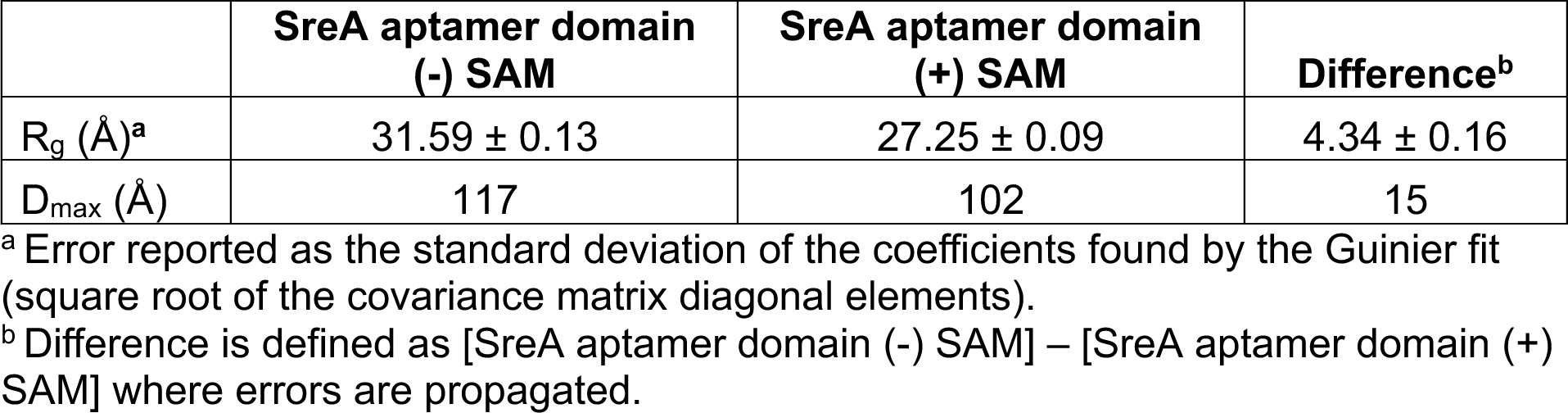
SAXS Data.

SreA aptamer domain adopts a secondary structure similar to other SAM-I riboswitches.

We used Switch-MaP^40^ to experimentally measure the SreA aptamer domain secondary structure in both the absence and presence of ligand (**Fig. 4A,B**). In both the SAM (+) and SAM (-) conditions, the SreA aptamer domain SHAPE-informed secondary structure is consistent with the conserved SAM-I secondary structure. The expected structure of four helices (P1-P4) arranged about the central junction is well supported by our SHAPE data.

**Figure 4.**
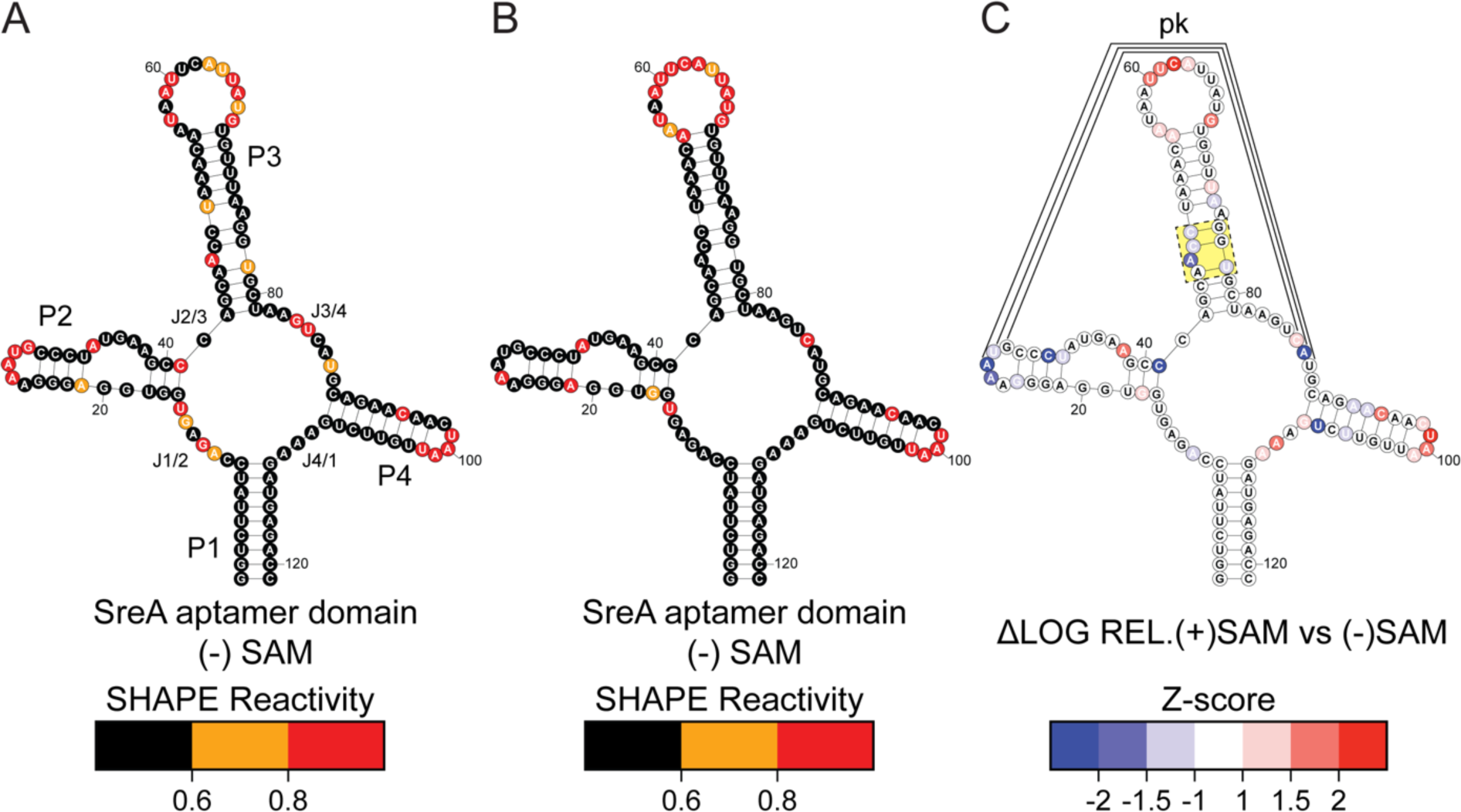
SreA aptamer domain adopts a SAM-I riboswitch secondary structure and undergoes SAM-dependent stabilization. **(A)** SreA aptamer domain secondary structure informed by SHAPE reactivity collected in the absence of SAM. Helical regions (P1-P4), junction regions (J1/2, J2/3, J3/4, and J4/1), and kink turn nucleotides (pk) are indicated. **(B)** SreA aptamer domain secondary structure informed by SHAPE reactivity collected in the presence of 1 mM SAM.**(C)** SreA aptamer domain secondary structure with Z-score analysis overlaid to indicate nucleotides that experienced SAM-dependent changes in SHAPE reactivity. Yellow box indicates nucleotides that experienced a SAM-dependent decrease in reactivity consistent with the expected mode of ligand binding. Plausible pseudoknot (pk) indicated by black lines.

To identify nucleotides that experienced SAM-dependent changes in reactivity, we applied an unbiased Z-score analysis comparing the SHAPE data from (+) SAM and (-) SAM conditions (**Fig. 4C**).^43^ Our Z-score analysis highlighted a number of significant differences in the SHAPE reactivities of the P2, P3, and P4 helices with SAM binding.

First, the P2 loop including A26, A27, and U28 experienced a large reduction in SHAPE reactivity in the presence of SAM. The conserved L2 – J3/4 pseudoknot is expected to include this loop, and is likely formed between A27:U88, U28:A87, and G29:C86. Notably, A87 is among the nucleotides with significantly reduced reactivity, supporting the formation of the expected pseudoknot. However, pseudoknot nucleotide C86 showed moderately increased reactivity upon SAM binding. Examination of reported ligand-bound SAM-I aptamer domain crystal structures reveals that the equivalent pseudoknot nucleotide is oriented with the 2’-OH solvent accessible and thus potentially hyper-reactive to acylation by a SHAPE reagent.^17,20^ The reactivities of the remaining pseudoknot nucleotides G29 and U88 while not among nucleotides with the most significant changes in SHAPE reactivity, did experience modest decreases in reactivity.

Second, the P3 nucleotides A47, C48, C49, and U78 also show significantly decreased reactivity in the presence of SAM. In reported crystal structures of ligand bound SAM-I riboswitches the highly conserved P3 asymmetric bulge forms an A-A_SAM_-U base triple with a dinucleotide platform further stabilizing the ligand through base stacking.^17,20^ The SAM dependent reduction of SreA aptamer domain SHAPE reactivity in this region support ligand binding through the previously reported mechanisms (**Fig. 4C** yellow box).^17,20^

Lastly, the apical loops of P3 and P4 show increased reactivity in the presence of SAM. Previous studies suggest that more stable helical regions can enable nucleotides in associated apical loops to more frequently sample unique orientations that are hyper-reactive to SHAPE reagents.^50^

SreA terminates transcription of the downstream genes.

To evaluate the transcriptional control of SreA on the downstream gene we performed single-round transcription termination assays. Here, we examined the extent to which an extended SreA RNA (containing the full SreA 5’ UTR and 30 nucleotides of the downstream gene) could terminate transcription in response to a range of SAM concentrations (1 nM - 8 mM) (**Fig. 5**). In the presence of 1 nM SAM, we observed that 34% of the transcription products were truncated, indicating that SreA can terminate transcription before the downstream gene is transcribed (**Table 3**). Increasing the concentration of SAM to 8 mM yielded the maximum amount of terminated transcription products (66%) and demonstrates the SAM dependence of SreA transcription termination (**Table 3**). The concentration of SAM required to induce half of this maximum termination response (T_50_) was 130 ± 10 nM (**Table 3**). During transcription with 130 nM SAM, the terminator and antiterminator conformations of the SreA expression platform are equally likely and the riboswitch permits transcription of equal amounts of full-length and terminated products. Therefore, the T_50_ value is an important sensitivity parameter that describes the minimum concentration of ligand required to “flip the switch” from mostly full-length transcription products to mostly terminated transcription products. Under the conditions tested and in a SAM-dependent manner, SreA downregulates the amount downstream gene transcribed by up to 32%.

**Figure 5.**
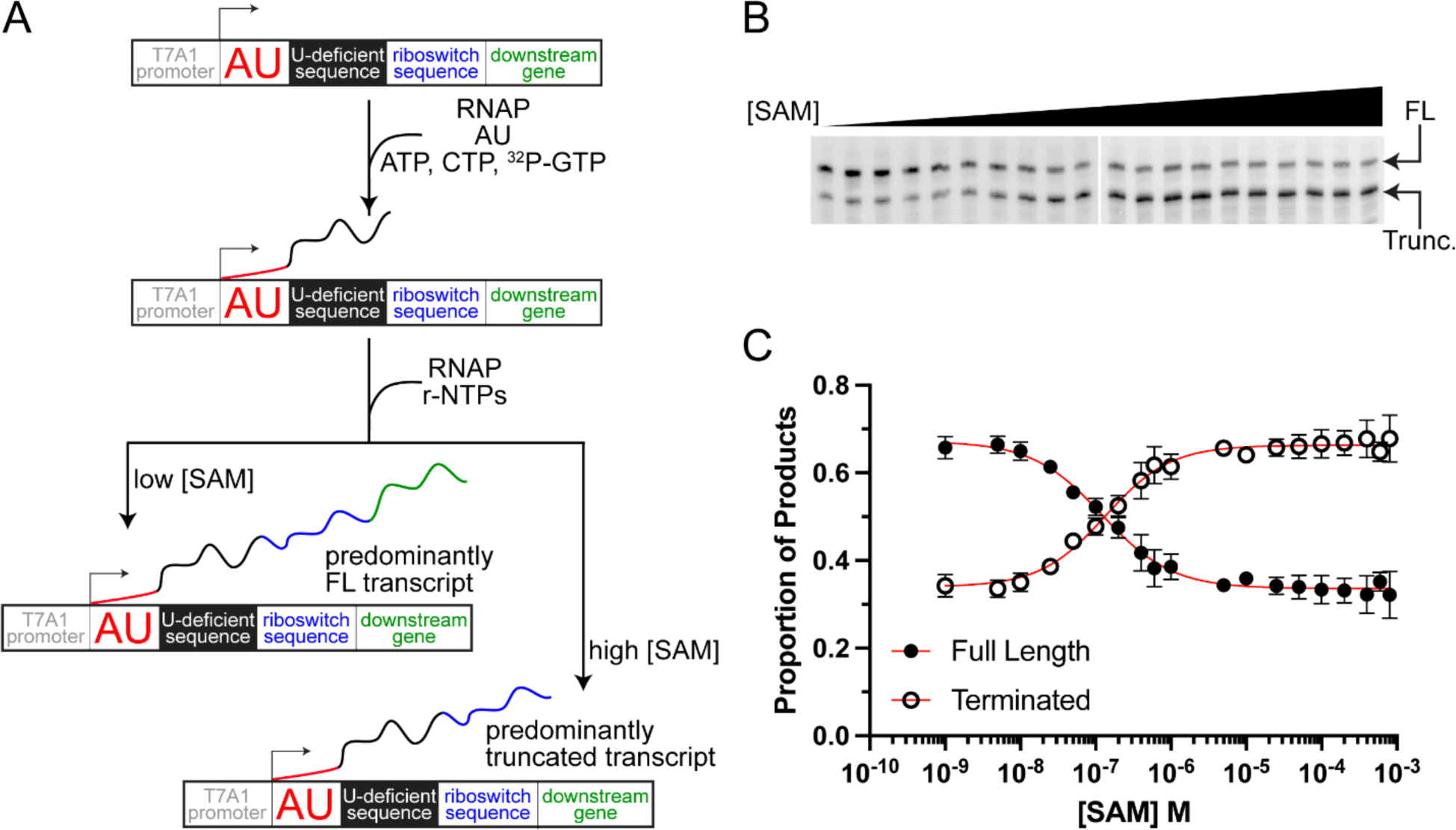
SreA controls the transcription of the downstream gene in a SAM-dependent manner. **(A)** Schematic overview of transcription termination assay. The halted RNA polymerase (RNAP) complex is formed by excluding UTP from the initial reaction mixture. ^32^P-GTP is incorporated into the RNA throughout the U-deficient sequence. The addition of a full set of NTPs allows the reaction to go to completion. Both full-length (FL) and truncated transcription products are formed in a SAM-dependent manner. **(B)** Phosphor imaging of PAGE gels used to resolve FL and truncated (trunc.) SreA transcription products formed with varying concentrations of SAM. **(C)** Transcription termination data collected in triplicate was fit to a four-parameter logistic curve to model the SAM-dependent transcriptional regulation by SreA.

**Table 3.**
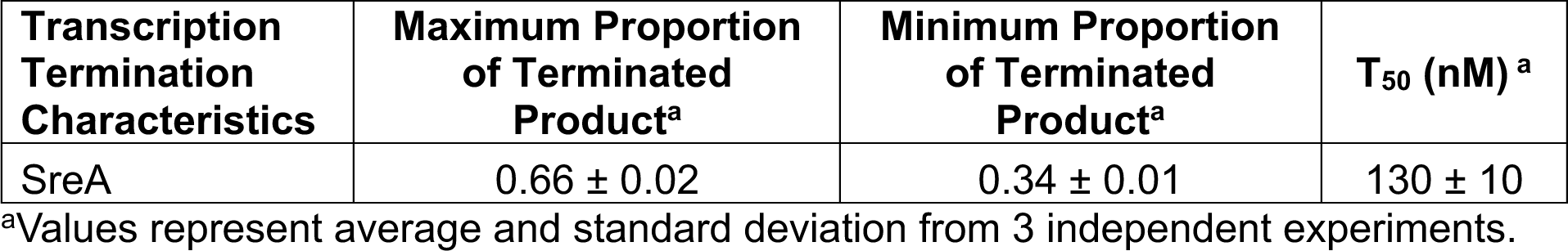
Transcription Termination Data.

## DISCUSSION

We validated SreA as a functional SAM-I riboswitch through characterization of ligand binding, structure, and transcriptional control. Our ITC experiments demonstrate that the SreA aptamer domain is competent to bind SAM and that it has similar binding properties to other known SAM-I riboswitches.^8,16,25,27^ The hepatocyte intracellular SAM concentration is 190 nM which is lower than the measured SAM binding K_D_ (490 nM) of SreA aptamer domain and that of many previously reported SAM-I riboswitches.^16,27,51^ The discrepancy between *in vitro* binding affinity and intracellular SAM concentration may be due, in part, to the truncation of the full-length riboswitch sequence and the stabilization of the aptamer domain construct. Additionally, the *in vitro* buffer conditions differ significantly from that of the cytoplasm of *L. monocytogenes*. It was recently shown that the metabolome of *E. coli* influences RNA stability.^52^ Likewise, it is reasonable to presume that the presence of *L. monocytogenes* metabolites, proteins, and other cytoplasmic components could influence SreA stability and ligand binding. It is also important to acknowledge that *in vivo* SAM binding occurs in a co-transcriptional context and the relevant binding affinity is not the equilibrium binding affinity determined in these assays.

The selectivity of the SreA aptamer domain for SAM is also similar to previously reported SAM-I riboswitches.^8,25^ SAH is toxic at high cellular concentrations and sulfur metabolism must be tightly regulated to prevent its accumulation.^53^ It is important for functional SAM-I riboswitches to be able to effectively discriminate between these ligands to properly regulate sulfur metabolism in the cell and prevent buildup of SAH.

SAM-I riboswitch aptamer domains adopt conserved and well-defined secondary and tertiary structures that are critical for ligand binding and transcriptional control. We used Switch-MaP and SAXS to characterize the structure of the SreA aptamer domain. Our chemical probing data support a secondary structure for the SreA aptamer domain that is consistent with the conserved SAM-I aptamer domain fold and inform on ligand-induced stabilization. Addition of 1 mM SAM decreased the SHAPE reactivity of SAM binding nucleotides in the P3 stem and of several nucleotides in the expected P2-J3/4 pseudoknot. The SAM-dependent changes in SHAPE reactivity validate that SreA binds SAM through the same highly conserved interactions as other known SAM-I riboswitches. Additionally, SAM binding compacted the SreA aptamer domain structure. SAXS has been used to characterize the extent of macromolecular compaction riboswitches undergo during folding and ligand binding.^21^ Magnesium is essential to the folding and compaction of SAM-I riboswitches and has been previously shown to largely pre-structure the aptamer domain into a ligand-binding competent state.^21^ SreA aptamer domain was similarly well folded in the presence of 5 mM MgCl_2_. Addition of 1 mM SAM further compacted the RNA and supports a previously described model where the conformationally flexible aptamer domain envelopes the ligand and buries it in the core of the RNA.^54^ During transcription the ligand induced conformational shift is thought to facilitate structural changes in the expression platform.^12,54^

Lastly, we demonstrated that full-length SreA can control the transcription of the downstream gene *in vitro*. Upon addition of 0-8 mM SAM, SreA increased transcription termination from 34% to 66% with T_50_ = 130 nM. Based on this result and reports that SAM is present at approximately 50 - 150 nM in human plasma, SreA is plausibly sensitive to *in vivo* SAM concentrations.^55–57^ One potential caveat to the results of the transcription termination experiments was that *E. coli* RNA polymerase was used rather than the native *L. monocytogenes* RNA polymerase. The *E. coli* enzyme has proficient promoter recognition and was previously reported to be an acceptable surrogate for transcription termination assays in *B.* subtilis.^18^ We evaluated the conservation of the *B. subtilis* and *L. monocytogenes* RNA polymerases to the *E. coli* RNA polymerase with BLAST, Jalview and the AACon webserver.^58–62^ The *B. subtilis* and *L. monocytogenes* RNA polymerases share a similar percent identity and extent of conservation with the *E. coli* RNA polymerase (**Fig. S1**) which supports our use of the *E. coli* RNA polymerase for SreA transcription termination assays.

In summary, we thoroughly validated SreA as a functional SAM-I riboswitch with a similar aptamer domain structure and ligand binding properties to previously characterized SAM-I riboswitches.^8,16–21^ Despite these similarities, full-length SreA controls the transcription of the downstream methionine ABC transporter genes in a distinct manner. SreA is highly sensitive to SAM but is a leaky riboswitch that permits transcription of the downstream genes during ligand scarcity and abundance. In contrast to the non-pathogenic bacteria *B. subtilis* and *T. tencongensis*, *L. monocytogenes* transitions from soil dwelling saprophyte to intracellular pathogen during a biphasic life cycle. The highly sensitive, but leaky transcription termination response of SreA may help ensure that genes for methionine import are transcribed even when the availability of environmental SAM changes during the *L. monocytogenes* life cycle. While SreA is not the only SAM-dependent riboswitch poised to control methionine import in *L. monocytogenes*, our data suggest that the methionine import genes following SreA would still be transcribed even when SAM is abundant in the environment.

In other bacteria, transcriptional control of methionine import genes can be more tightly regulated. For comparison, the transcription of methionine ABC transporter genes in *B. subtilis* are also under the control of a SAM-I riboswitch. The *yusC* SAM-I riboswitch is sensitive to μM SAM concentrations and increases transcription termination from 52% to 90% at high concentrations of SAM.^16^ This means that when SAM is abundant in the environment, genes for methionine import are rarely transcribed. Interestingly the *cysH* SAM-I riboswitch from *B. subtilis* had a similar maximum percent transcription termination relative to SreA (63% and 66%, respectively) but controls transcription of a biosynthetic gene rather than a transport gene.^16^ Together these observations highlight how bacteria can use one class of riboswitch to achieve distinct regulatory outcomes.

Because the SAM-I riboswitch aptamer domain is highly conserved, it is possible that differences in riboswitch function may be due in part to differences in the poorly conserved expression platforms. In fact, a study focused on the *Clostridium beijerinckii* ZTP riboswitch demonstrated that variations in the expression platform sequence could tune the transcription termination response to a high dynamic range by imposing different sizes of kinetic barriers to the strand displacement mechanism required for regulation.^63^ Strand displacement has emerged as a ubiquitous mechanism that guides the conformational rearrangements of transcriptional riboswitches.^14,15,44,64–67^ For SAM-I riboswitches, ligand binding in the aptamer domain likely represses a strand displacement mechanism between P1 stem and the terminator hairpin in the expression platform that would otherwise form the antiterminator hairpin. Though we did not characterize the expression platform in the current study, we did analyze both P1 stem stability and maximum percent terminated product for SreA (this work) and the *B. subtilis* SAM-I riboswitches which were characterized using similar *in vitro* transcription termination assays.^16^ This analysis revealed a trend (R^2^ = 0.77) where greater P1 stem stability is correlated with greater terminated product formation (**Fig. S2**). This correlation supports a model in which expression platform strand displacement may underly the differences observed SAM-I riboswitch transcription termination responses.

### Data Availability

All plasmids and raw data files generated in this work are available upon request. Experimental SAXS data of the SreA aptamer domain have been deposited in the Small Angle Scattering Biological Data Bank under accession codes SASDSK9 (-SAM) and SASDSL9 (+SAM). Raw sequencing data and associated processed data files were deposited in GEO (GSE266872).

## Acknowledgments

We acknowledge the funding support from the National Science Foundation (MCB-1942398 to S.C.K.). C.A.W is supported by an NCI transition award (K22CA262349) and startup funds from the University of Michigan Center for RNA Biomedicine and an NCI Support Grant for the Rogel Cancer Center (P30CA046592). We thank Dr. Nils Walter and Dr. Adrien Chauvier for their assistance with transcription termination construct design and methods. We gratefully acknowledge the Advanced Photon Source, Dr. Jesse Hopkins, Dr. Maxwell Watkins, and the staff of BioCAT. This research used resources of the Advanced Photon Source, a U.S. Department of Energy (DOE) Office of Science User Facility operated for the DOE Office of Science by Argonne National Laboratory under Contract No. DE-AC02-06CH11357. This project was supported by grant P30 GM138395 from the National Institute of General Medical Sciences of the National Institutes of Health. Use of the Pilatus 3 1M detector was provided by grant 1S10OD018090-01 from NIGMS. The content is solely the responsibility of the authors and does not necessarily reflect the official views of the National Institute of General Medical Sciences or the National Institutes of Health.

