## Supplementary Data for "Functional Validation of SAM Riboswitch Element A from *Listeria monocytogenes*"

**Table S1. Riboswitch Sequences**

| **Riboswitch** | **Sequence 5′-3′** |
| --- | --- |
| SreA SAM-I riboswitch | caagaaacgucacuauagucacaaaugaagugauuuaauaauauguagaaaucuuauccagagugguggagggaaaugcccuaugaagcccagcaaccuaaacaauaauucauuauguguuuaaggugcuaagucaugcagaacaacuaauuguucugaaagaugagaaggaaguuaguccauuugaaaaaaugcugccuuucugcucaucagcagaaaggcuuuuuuu |
| *yitJ* SAM-I riboswitch | ACUUCCUGACACGAAAAUUUCAUAUCCGUUCUUAUCAAGAGAAGCAGAGGGACUGGCCCGACGAAGCUUCAGCAACCGGUGUAAUGGCGAUCAGCCAUGACCAAGGUGCUAAAUCCAGCAAGCUCGAACAGCUUGGAAGAUAAGAAGAGACAAAAUCACUGACAAAGUCUUCUUCUUAAGAGGACUUUUUUUAUUU |

**Table S2. SreA Constructs**

| **Construct Name** | **Sequence 5′-3′** |
| --- | --- |
| SreA gene block | AGTTACGAATTCTAATACGACTCACTATAGGGATCGACGTTTCTTGCTGATGAGTCCGTGAGGACGAAACGGTACCCGGTACCGTCCAAGAAACGTCACTATAGTCACAAATGAAGTGATTTAATAATATGTAGAAATCTTATCCAGAGTGGTGGAGGGAAATGCCCTATGAAGCCCAGCAACCTAAACAATAATTCATTATGTGTTTAAGGTGCTAAGTCATGCAGAACAACTAATTGTTCTGAAAGATGAGAAGGAAGTTAGTCCATTTGAAAAAATGCTGCCTTTCTGCTCATCAGCAGAAAGGCTTTTTTTGGATCCCTAGATA |
| SreA transcription termination gene block | AGTTACGAGCTCTCCAGATCCCGAAAATTTATCAAAAAGAGTATTGACTTAAAGTCTAACCTATAGGATACTTACAGCCATCACGGGACACCCAACCAAGAAACGTCACTATAGTCACAAATGAAGTGATTTAATAATATGTAGAAATCTTATCCAGAGTGGTGGAGGGAAATGCCCTATGAAGCCCAGCAACCTAAACAATAATTCATTATGTGTTTAAGGTGCTAAGTCATGCAGAACAACTAATTGTTCTGAAAGATGAGAAGGAAGTTAGTCCATTTGAAAAAATGCTGCCTTTCTGCTCATCAGCAGAAAGGCTTTTTTTGTATATCAGAATGTAGAAAAGGTGATAGAGAAGCTTCTAGAT |

**Table S3. Oligo Sequences**

| **Oligo Name** | **Sequence 5′-3′** |
| --- | --- |
| SreA AD Fwd | AATAGTGAATTCTAATACGACTCACTATAGGTCTTATCCAGAGTGGTGGAGGGAAATGCC |
| SreA AD Rev^a^ | mGmGTCTCATCTTTCAGAACAATTAGTTGTTCTGCATGACTTAGCACCTT |
| SHAPE 5′ Adapter^b,c^ | rAppNNNNNNNCTGTCTCTTATACACATCTCCGAGCCCACGAGAddC |
| SHAPE TSO^d^ | TiMe-isodCGTCGGCAGCGTCAGATGTGTATAAGAGACAGNNNNNNNrGrGrG |
| RT Primer | GTCTCGTGGGCTCGG |
| i5 multiplexing primer | AATGATACGGCGACCACCGAGATCTACAC-X_8_-TCGTCGGCAGCGTC |
| i7 multiplexing primer | CAAGCAGAAGACGGCATACGAGAT-X_9_-GTCTCGTGGGCTCGG |

^a^ m denotes a 2′-O-methyl modification.

^b^ rApp denotes an adenylate group.

^c^ ddC denotes a dideoxy cytosine.

^d^ Me-isodC denotes a 5-methyl isodeoxycytosine.

^e^ X denotes variable nucleotides used for unique molecular identifiers during sequencing.

**Table S4. SEC-SAXS data acquisition, sample details, data analysis, model fitting, and software used.**

| **(*a*) Sample details** | | |
| --- | --- | --- |
| Organism | *Listeria monocytogenes* | *Listeria monocytogenes* |
| Source | *In vitro* transcribed RNA | *In vitro* transcribed RNA |
| DNA/RNA(s)^a^ | URS000028C263_1288293 | URS000028C263_1288293 |
| Ligand | N/A | *S*-adenosyl-L-methionine (1mM) |
| Stoichiometry of components | N/A | 1:1 |
| Solvent composition | 50 mM Potassium Phosphate pH 6.5, 135 mM NaCl, 5 mM MgCl_2_ | 50 mM Potassium Phosphate pH 6.5, 135 mM NaCl, 5 mM MgCl_2_ |
| Sample temperature (°C) | 22 | 22 |
| In beam sample cell | 1.0 mm ID quartz capillary | 1.0 mm ID quartz capillary |
| Sample injection concentration, mg/ml | 2.5 | 2.5 |
| Sample injection volume, mL | 0.200 | 0.200 |
| SEC column type | Superdex 200 10/300 Increase | Superdex 200 10/300 Increase |
| SEC flowrate, mL/min | 0.6 | 0.6 |
| **(*b*) SAS data collection** | | |
| Data acquisition/reduction software | SEC-SAXS. Size separation used a Superdex 200 Increase 10/300 GL column and a 1260 Infinity II HPLC (Agilent Technologies). UV data was measured in the Agilent. SAXS data was measured in a sheath-flow cell (Kirby et al., 2016), effective path length 0.542 mm. Data reduced with RAW 2.1.3.^1^ | SEC-SAXS. Size separation used a Superdex 200 Increase 10/300 GL column and a 1260 Infinity II HPLC (Agilent Technologies). UV data was measured in the Agilent. SAXS data was measured in a sheath-flow cell (Kirby et al., 2016), effective path length 0.542 mm. Data reduced with RAW 2.1.3.^1^ |
| Source/instrument description or reference | BioCAT facility at the Advanced Photon Source beamline 18ID with Eiger2 XE 9M (dectris) detector | BioCAT facility at the Advanced Photon Source beamline 18ID with Eiger2 XE 9M (dectris) detector |
| Measured *q*-range  (*q_min_* – *q_max_*; Å^-1^) | 0.0027 to 0.42 Å^-1^ | 0.0027 to 0.42 Å^-1^ |
| Method for scaling intensities | Glassy Carbon, NIST SRM 3600 | Glassy Carbon, NIST SRM 3600 |
| Exposure time(s), number of exposures. For SEC-SAS, final number of sample frames used for averaging. | 0.5 s exposure time with a 1 s total exposure period (0.5 s on, 0.5 s off). 2771 exposures with 15 frames used for averaging. | 0.5 s exposure time with a 1 s total exposure period (0.5 s on, 0.5 s off). 3000 exposures with 12 frames used for averaging. |
| **(*c*) SAS-derived structural parameters** | | |
| Methods/Software | Radial averaging; frame comparison, averaging, and subtraction done using BioXTAS RAW 2.1.3^1^ | Radial averaging; frame comparison, averaging, and subtraction done using BioXTAS RAW 2.1.3^1^ |
| **Guinier Analysis** | | |
| I(0) ± σ (a.u) | 0.01 ± 2.26e-5 | 0.01 ± 2.1e-5 |
| R_g_ ± σ (Å) | 31.59 ± 0.13 | 27.25 ± 0.09 |
| min < qR_g_ < max limit | 0.08 < qR_g_ < 1.3 | 0.07 < qR_g_ <1.33 |
| Linear fit assessment (r^2^) | 0.96 | 0.98 |
| **P(r) analysis** | | |
| I(0) ± σ (a.u.) | 0.01 ± 2.25e-5 | 0.01 ± 2.19e-5 |
| R_g_ ± σ (Å) | 32.45 ± 0.13 | 28.01 ± 0.11 |
| D_max_ (Å) | 117.0 | 102.0 |
| q-range (Å^-1^) | 0.003 – 0.253 | 0.003 – 0.294 |
| P(r) fit assessment (χ^2^) | 0.882 | 0.976 |
| **(*d*) Scattering particle size** | | |
| Methods/Software | BioXTAS RAW 2.1.3^1^ | BioXTAS RAW 2.1.3^1^ |
| Porod volume, *V_p_* (Å^3^) | 97500 | 81200 |
| Molecular weight estimates from chemical composition (kDa) | 39.22 | 39.22 |
| Molecular weight estimates from SAS, concentration independent method (V_c_) (kDa) | 44.1 | 40.1 |
| **(e)** **Data and model deposition** | | |
| SASBDB IDs | SASDSK9 | SASDSL9 |

^a^ RNACentral (<https://rnacentral.org/>) ascension numbers are provided for the entry “*Listeria monocytogenes* QOC1 SAM”. Relative to the RNAcentral entry, the SreA aptamer domain sequence used in this study has A1 substituted by two G residues and A119 substituted by two C residues.

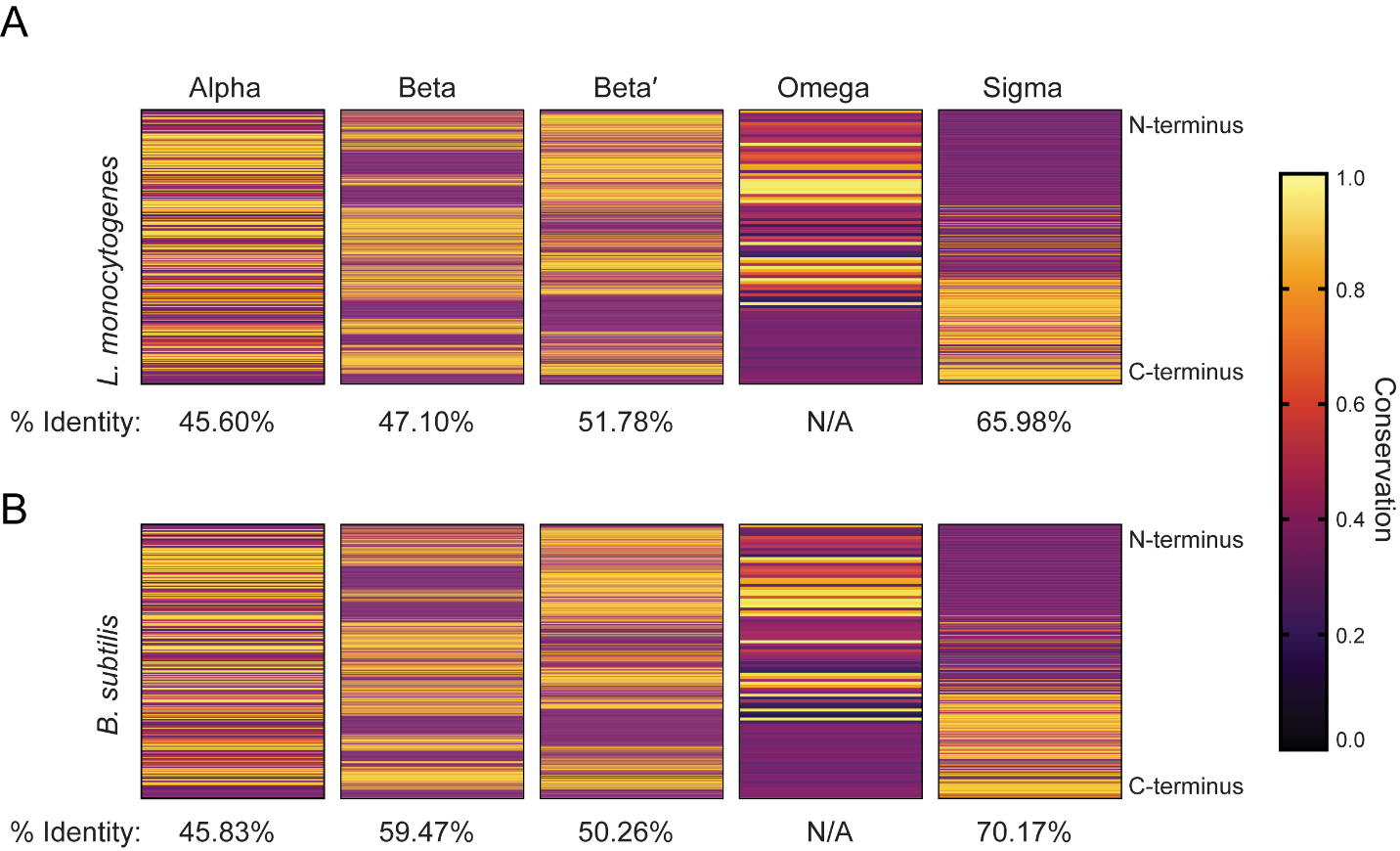

**Figure S1. B. subtilis and L. monocytogenes RNA polymerasess have similar conservation to the E. coli RNA polymerase.** Percent identities determined by BLAST and N/A indicates no significant percent identity.^2^ 17 conservation metrics from the AACons webserver were normalized and averaged to calculate the amino acid conservation scores.^3–5^ **(A)** Comparison of the alpha, beta, beta**′**, and omega subunits L. monocytogenes RNA polymerases as well as sigma factor 70 to the equivalent E. coli proteins used in the transcription termination assays. **(B)** Comparison of the alpha, beta, beta ′, and omega subunits B. subtilis RNA polymerases as well as sigma factor 70 to the equivalent E. coli proteins used in the transcription termination assays.

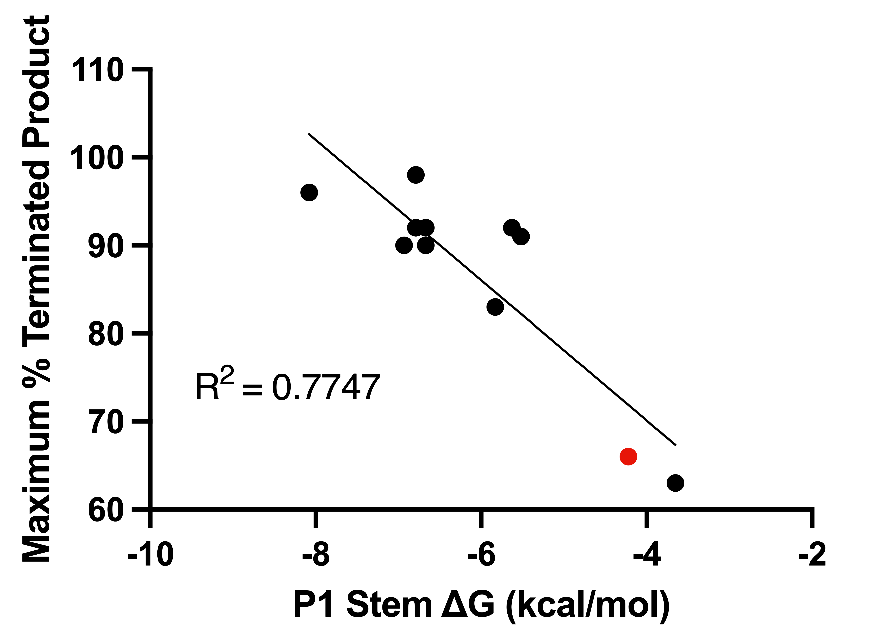

**Figure S2. P1 stem stability and extent of transcription termination.** The folding stability (ΔG) of P1 stem sequences from SreA and the Bacillus subtilis SAM-I riboswitches (ykrT/mtnK, yxjG, yitJ/samT, yjcI/metI, metE, yusC/metN, yoaD, ykrW/mtnW, yxjH, and cysH) were calculated using nearest neighbor rules. Folding stabilities for the B. subtilis riboswitches were plotted against previously reported maximum percent terminated products (black data point).^6^ SreA folding stability was plotted against the maximum percent terminated product from the present study (red data point). P1 folding stability is negatively correlated with maximum percent terminated product (R^2^ = 0.7747).
